# Graphene Oxide reinforced Agarose-Hydroxyapatite Bioprinted 3-D Scaffolds for Bone Regeneration

**DOI:** 10.1101/2021.12.24.474115

**Authors:** Umakant Yadav, Shiva Kumar

**Author notes:** Corresponding author, (Dr. Umakant Yadav).

## Abstract

Three-dimensional (3D) bioprinting is an emerging technology for fabricating cells, biomaterials and extracellular matrix (ECM) into customized shapes and patterns. Here, we report additive manufacturing to create a customized 3D bioactive constructs for regenerative medicine. We have attempted to emphasize the use of agarose and graphene oxide as a promising material for the conceptualization of bioink unpaid to its unique physicochemical properties. The 3D printed structure is able to regenerating bone tissues and regulates the cellular differentiation without any significant morphological changes. The presence of graphene oxide enhances the osteoinductive behavior of the developed scaffolds, which is further supplemented by encapsulating human mesenchymal stem cells (hMSCs) on the 3D printed scaffolds.

A significant enhanced expression of early osteogenic markers like morphogenetic protein (BMP), Runx-2, collagen-1, osteopontin, osteocalcin as well as mineralized ECM are observed on agarose-hydroxyapatite and graphene oxide 3D printed scaffolds compared to agarose-hydroxyapatite 3D printed scaffolds. Thus, the outcomes of the developed 3D bioprinted scaffolds provide a promising strategy for development of personalized bone grafts for tissue regeneration.

## 1. INTRODUCTION

Currently, bone disorders have become a major problem for the present era due to increased burden on life style and trauma in our population. Approximately 200 000 incidences of bone (CMF) injuries occur annually due to trauma, congenital malformations, and arthroplasty interventions [1]. The major approach for tissue engineering has attempted to develop a suitable scaffold that mimic the natural extra cellular matrix and enhances the bone repair by suitable microenvironment for osteoblast differentiation [2]. The major role of regenerative medicine is to develop personalized implants that restore tissue functionality without any obvious toxic effects. Three-dimensional scaffolds can enhance bone regeneration by representing suitable microenvironment that facilitates incursion of cells from neighboring tissues, proliferation, differentiation, development of bone extracellular matrix (ECM) and vascular beds [3]. Ceramic based materials has been extensively used in bone tissue engineering, due to their high mechanical strength, biocompatibility and osteo-conductivity, apart from this degradation, and fragile mechanical properties are important limits to be considered ideal material for clinical applications [4]. Natural polymer based materials has been used in tissue engineering due to their biodegradation properties as well as excellent cell compatibility [5]. Due to unique physicochemical properties, minimum immune inflammatory response and good biocompatiblilty agarose has become most promising polymeric material in the field of tissue engineering [6]. In the last decay, graphene oxide (GO) has gain more attention in the field of regenerative medicine due to its unique physico-chemical, mechanical and biocompatible nature [7]. Human mesenchymal stem cells (hMSCs) have become an attractive candidate in regenerative medicine for bone repair due to its osteo stimulatory properties [8]. To the best of our knowledge, no reports have been available to evaluate the cellular differentiation and osteogenicity of 3D printed agarose-hydroxyapatite and graphene oxide scaffolds.

The objective of the current study is to develop 3D printed agarose-hydroxyapatite and graphene oxide scaffolds, encapsulation of hMSCs and in vitro osteogenic differentiation. The osteogenic differentiation of hMSCs were analyzed by confocal microscopy and RT-PCR assays.

## 2. MATERIALS AND METHODS

### 2.1. Materials

Calcium hydroxide (Ca(OH)2), orthophosphoric acid (H3PO4) and glacial acetic acid (CH3COOH) were procured from Sigma-Aldrich. Dulbecco’s Modified Eagle’s Medium (DMEM) high-glucose media, Fetal bovine serum (FBS) and trypsin-ethylenediaminetetraacetic acid (Trypsin-EDTA) were from Thermo Fisher Scientific (Massachusetts, U.S.A.). hMSCs cells were procured from National Centre for Cell Science (NCCS, Pune, India). 3-(4,5-Dimethylthiazol-2-yl)-2,5-diphenyltetrazolium bromide (MTT) and agarose were purchased from Himedia Laboratories Pvt (SRL, India).

### 2.2. Synthesis of hydroxyapatite (HAP)

HAP was synthesized by a simple chemical precipitation method [10]. In short, 1.85 gram of calcium hydroxide[Ca(OH)2,0.1 M Ca^2+^] was dissolved in 1% glacial acetic acid[CH3COOH] and stirred for 2 h at 60°C.In anotherflask,0.1 M orthophosphoric acid [H3PO4] solution was prepared in dilute ammonia and pH was maintained~11-12.These solutions were mixed together with a constant rate (4.0 mLmin^-1^) and stirred vigorously for 12 h. The formed precipitate was allowed to settle followed by washing thrice with double distilled water with a series of intermittent incubation on water bath at 80°C.Finally, the precipitates were subjected to centrifugation at 4000 rpm for 5 min to obtain consolidated hydroxyapatite.

### 2.3. Synthesis of Graphene oxide (GO)

GO have been synthesized by improved Hummers method with a slight modification [9]. In brief, 0.0015 kg of graphite powder have been pre-oxidized by reacting it with a mixture of 40 ml of 98% H2SO_4_, 5g K_2_S2O_8_ and 5 g of P_2_O_5_ for 4h at 80°C. Further oxidation have been achieved by adding the pre-oxidized graphite to a mixture of concentrated H2SO_4_-H_3_PO_4_ (v/v: 180: 13) with constant stirring. After 5 min, 9 g of KMnO_4_ have been added to the mixture and the stirring is continued for 15h at 50°C. The reaction is stopped and the reactants were allowed to cool at RT followed by pouring of 200 ml of ice and 1.5 ml of H_2_O_2_ (30%). Multiple washings of the material have been carried out with DI water, 30% HCl and ethanol and finally coagulated with ether. The obtained semi-solid material has been vacuum dried overnight to obtain brown graphene oxide (GO) powder.

### 2.4. Bioink Formulation

Agarose-hydroxyapatite and graphene oxide was synthesized according to the protocol reported by Chimene, D. et al. [11]. In brief, 1.5 g of agarose was dissolved into 100 mL of 1 × phosphate buffered saline (PBS) by heating at 90° C. After complete dissolution 1.0% (w/v) solution of graphene oxide and 4.0% (w/v) solution of hydroxyapatite were mixed in agarose solution, and vigorously stirred for 4 h at 90 ° C. After that, the solution was preserved at 40 ° C for 12 h.

### 2.5. 3D Printing

Agarose-hydroxyapatite and graphene oxide constructs has been fabricated using HYREL System 30M 3D printer. Preserved solution was laden into a HYREL VOL-25 extruder (HYREL L.L.C., Norcross, GA) equipped with a adapter and 23 gauge stainless steel needle. Once connected to the printer, constructs were modeled in Solid works 3D CAD Design, exported as an STL file, and imported into Slic3r version 1.2.9. Overall, this process converts the Solid works design into layer-by-layer instructions for the printer, or G-code. The G-code files are then imported into HYREL’s proprietary software (Repetrel Rev2.828) and printed at room temperature onto glass slides. Repetier Host was used to control the 3D printer. All printed constructs were programmed to have a layer height of 200 μm and a speed of 0.20 mm s - 1.

### 2.6. Haemolytic analysis

Haemolytic activities of developed scaffolds were evaluated by detecting the haemoglobin release from human red blood cells [12]. Healthy goat blood was collected from local butcher’s shop, Kanpur, India. The blood was centrifuged at 3000 rpm at 4°C for 10 min and washed thrice with PBS solution (pH 7.4). Red blood cells (RBCs) were obtained and the previous volume was maintained using PBS solution. Then, the RBC suspension was diluted ten times to a concentration of 10% (v/v) with PBS. Furthermore, 1.5 mL of the RBC suspension was taken in autoclaved 2.0 mL centrifuge tubes and incubated scaffolds for 2 h. Afterward, all the suspensions were centrifuged at 3000 rpm at 4°C for 10 min, and the absorbance of supernatant was recorded at 540 nm. Triton-X was taken as negative control (100% haemolysis) and PBS as positive control (0.0% haemolysis). The percentage haemolysis was expressed by the formula given below.

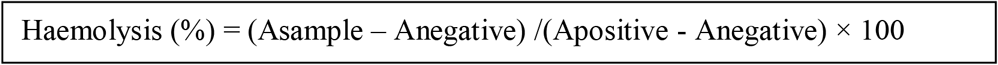

### 2.7. Cell seeding on 3D Printed Scaffolds

Before cell seeding all the scaffolds were sterilized by three times washing with 70% ethanol followed by UV sterilization in a 6 well culture plates. After sterilization scaffolds were dipped in Dulbecco’s Modified Eagle Medium (DMEM)), supplemented with 10% fetal bovine serum (FBS), and 1% penicillin/streptomycin under controlled atmospheric conditions (37°C temperature and 5% CO2) for 12 h. Excess medium has been removed from the scaffolds before cell seeding. In brief, hMSCs cells were seeded in 6 well culture plates with a density 1×10^6^cells/well and incubated for 24 h allowing for initial attachment over scaffolds. After 24 h, old media was replaced with fresh medium for 16 days. The scaffolds without graphene consider as control.

### 2.8. Cell Viability

Viability was analyzed through a live–dead assay. Specifically, cells were stained with a solution of ethidium homodimer (4 μL/mL) and calcein AM (2 μL/mL) in PBS for 30 min, and rinsed with PBS. The cell morphology was observed by fluorescence microscopy (Leica DMi Microscope).

### 2.9. Alkaline phosphatase Activity

Alkaline phosphatase (ALP) activity was measured using colorimetric ALP assay kit (Beacon, India). In brief, cell culture supernatant samples collected during the *in vitro* differentiation process, following manufacturer instructions. All experiments were performed in triplicate.

### 2.10. DNA Content Analysis

To evaluate the cell compatibility of fabricated 3D scaffolds, hMSCs proliferation was evaluated in the form of total DNA concentration. In brief, total DNA content was isolated from the 3D constructs at 14 and 21 days using a DNA extraction kit (Sigma). The isolated DNA contents were collected and quantified by using Nano drop (Thermo Scientific-2000).

### 2.11. Osteogenic markers expression

The hMSCs were seeded on different scaffold has been trypsinized and collected on 8^th^ and 21^st^ days of culture, day. Total RNA was extracted using chloroform-ethanol precipitation method with TRIzol reagent. RNA pellets were washed three times with 70% ethanol; air-dried and dissolves in DEPC-treated water. After DNase I treatment RNA was quantified with Nanodrop to test for integrity. After that, 2 μg of RNA have been used to synthesize first strand cDNA using iScript cDNA Synthesis kit. The ECM protein-coding gene expression like aggrecan core protein (ACAN), collagen-1A2 (COL1A2), Cartilage oligomeric matrix protein (COMP), collagen-1A1 (COL1A1), collagen-3A1 (COL3A1), bone morphogenetic protein-1 (BMP1), bone morphogenetic protein-6 (BMP6) and runt related transcription factor-2 (RUNX2) were evaluated using real-time quantitative polymerase chain reaction (qRT-PCR).

### 2.12. Statistical Analysis

All data obtained from experiments were processed using SPSS 16.0 and represented as mean ± standard deviation (SD). Statistical comparisons of all data were performed by one-way analysis of variance (ANOVA), followed by Dunnet post hoc tests. P values < 0.05 were considered to indicate a statistically significant difference.

## 3. Result and discussion

The main objective of the present study was to develop 3D bioprinted scaffolds and in vitro evaluation of osteogenic differentiation of hMSCs in articular chondrogenic lineage.

### 3.1. Fabrication of 3D bioprint

Effective bioinks for 3D bioprinting by additive methods requires ink flow through a needle (printability) to display shear-thinning properties, allowing the bioink to freely flow from the printing nozzle in a way that reduces the shear stresses. The bioink should rapidly recuperate its stickiness after extrusion in order to minimize further shape deformities.

Agarose used in the present study, has assure as a material for bioprinting due to its excellent biocompatibility, ease in forming gels, and shear thinning properties [13].

Furthermore, the high and rough surface area of hydroxyapatite can also be enhanced for drug-delivery application, promoting sustained and prolonged release of therapeutics encapsulated in to the 3D printed scaffolds [14]. Here, here we have printed a simple rectangle with a 9.0 mm length and 0.26 mm thick for investigating cell-scaffold interactions. Hermenean et al., 2017 reported that fabrication of graphene oxide in 3D scaffolds accelerates bone regeneration in a critical-size mouse calvarial defects model [15]. In this study, we hypothesized that encapsulation of hMSCs on the surface of 3D-printed scaffolds will promote osteogenesis and enhances bone healing capabilities.

### 3.2. Haemolysis assay

The hemocompatibility of any nanomaterial with human blood components is a crucial toxicological consideration for the successful application of nanomaterials in biomedical applications. Blood is mainly composed of blood cells and blood plasma, blood cells like, RBCs, WBCs and platelets that are suspended in blood plasma. To address blood compatibility, haemolysis assay was also performed against different scaffolds with RBCs, and results are presented in Figure 5. Triton-X was taken as positive control (100% haemolysis) and PBS as negative control (0.0% haemolysis). The haemolysis percentage of RBCs exposed to different scaffolds was negligible (less than 1.0%). Hence, it can be inferred that these scaffolds are an excellent compatible material and can be used for the biomedical applications.

### 3.3. Cell Viability

Biocompatibility of the fabricated 3D scaffolds was evaluated by the live/dead assay (Figure 3). From the fluorescence signals, it has been observed that hMSCs were uniformly distributed over the 3D bioprinted scaffolds. On 14^th^ day, the percentage of viable hMSCs increased (87.87%) significantly as compare to 7^th^ day. It has been reported that hydroxyapatite based scaffolds are bioresorbable and supports cellular components of newly formed bone [16]. From this, we can conclude that live/dead staining confirmed that the developed 3D scaffolds supports cell viability.

### 3.4. DNA Content Analysis

In vitro cell proliferation over 14 days was measured as quantitative evidence by amount of DNA (Figure 2b). Obtained data clearly reveals that incorporation of graphene oxide enhances the amount of DNA on the scaffolds. A significant increase in DNA concentration was observed for agarose-hydroxyapatite-graphene oxide scaffold. This suggests that agarose-hydroxyapatite-graphene oxide scaffolds support the optimal cell adhesion and proliferation of hMSCs without any obvious toxicity. Therefore, it is essential for this study to investigate the ability of agarose-hydroxyapatite-graphene oxide scaffolds to support bone and vascular intrusion, which finally leads to bone construction and the vasculogenesis.

**Figure 1.**
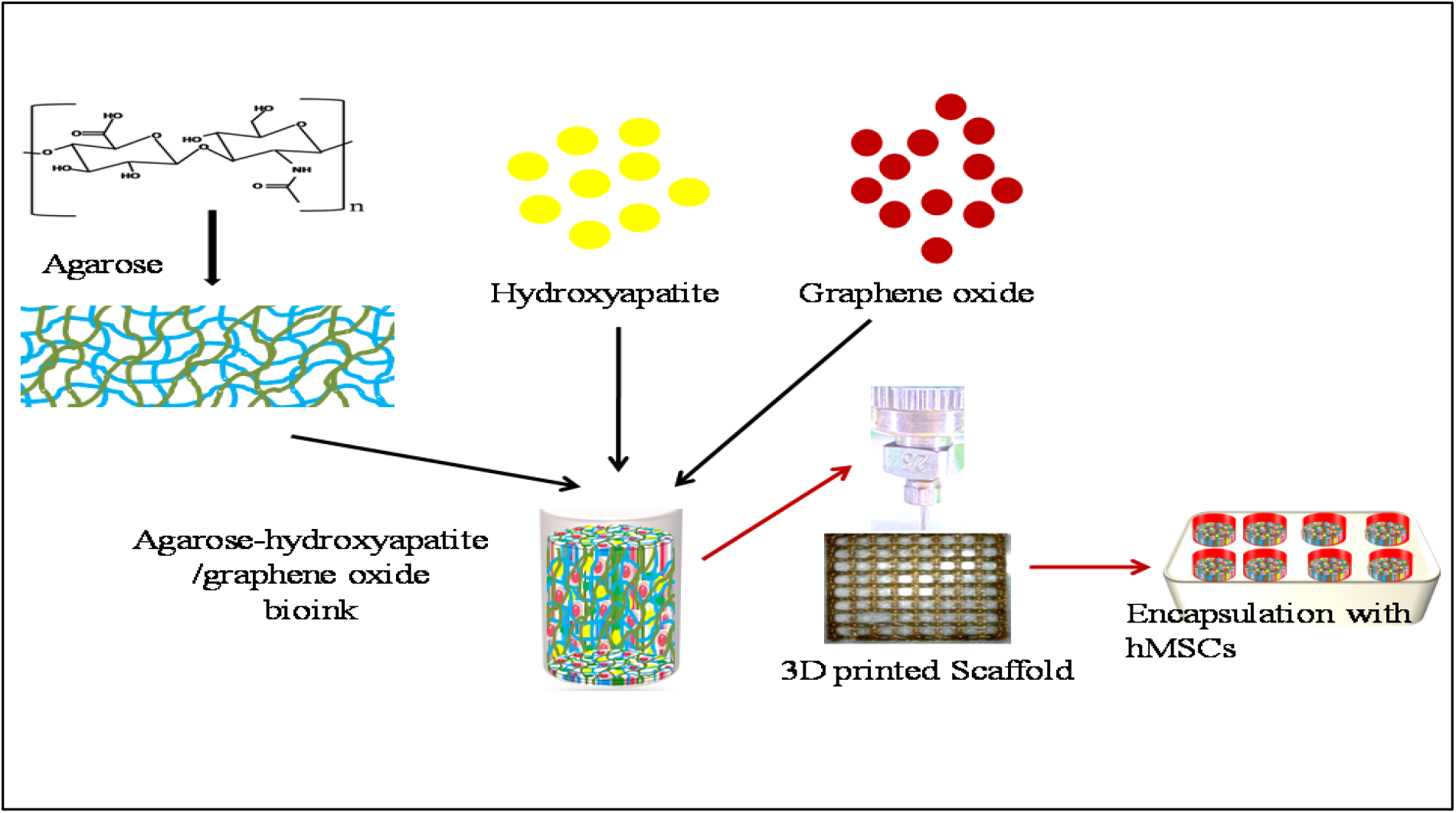
Schematic illustration of bioink preparation, printing of 3D scaffolds and hMSCs encapsulation.

**Figure 2.**
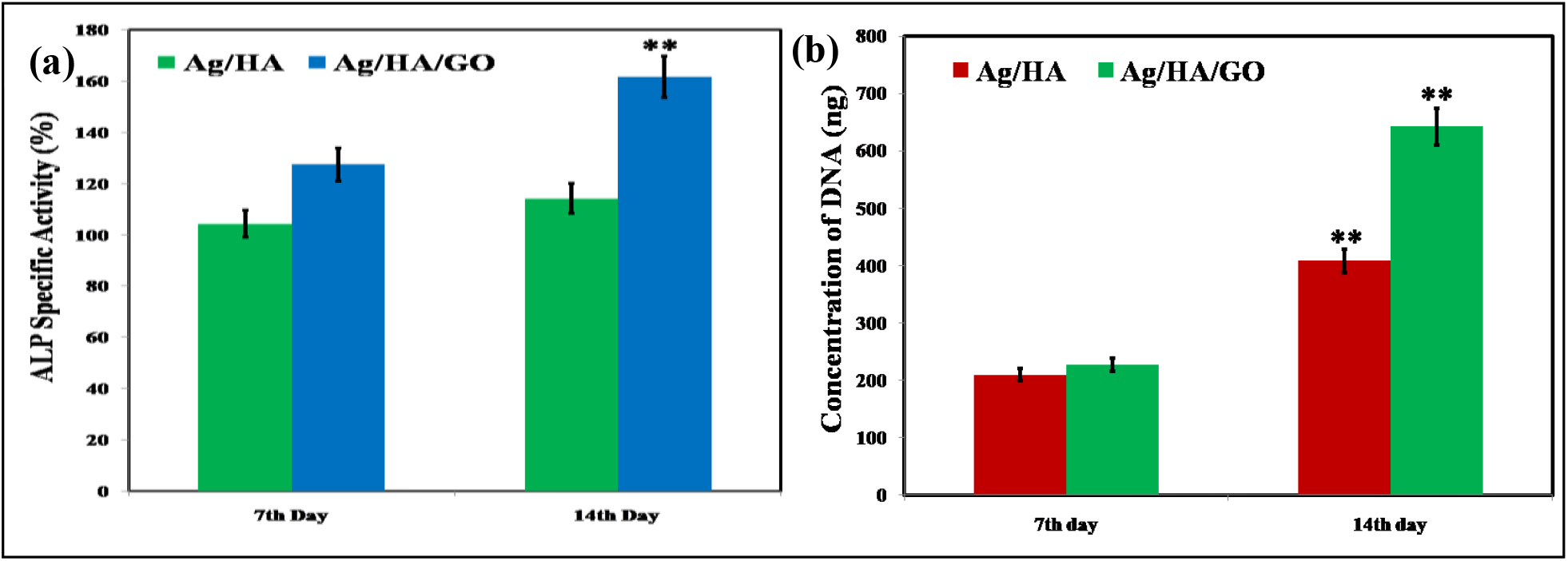
Showing the scaffolds biocompatibility in terms of alkaline phosphatase activity (a), and DNA concentration (b). Data shows Mean ± SD, n = 3.

**Figure 3.**
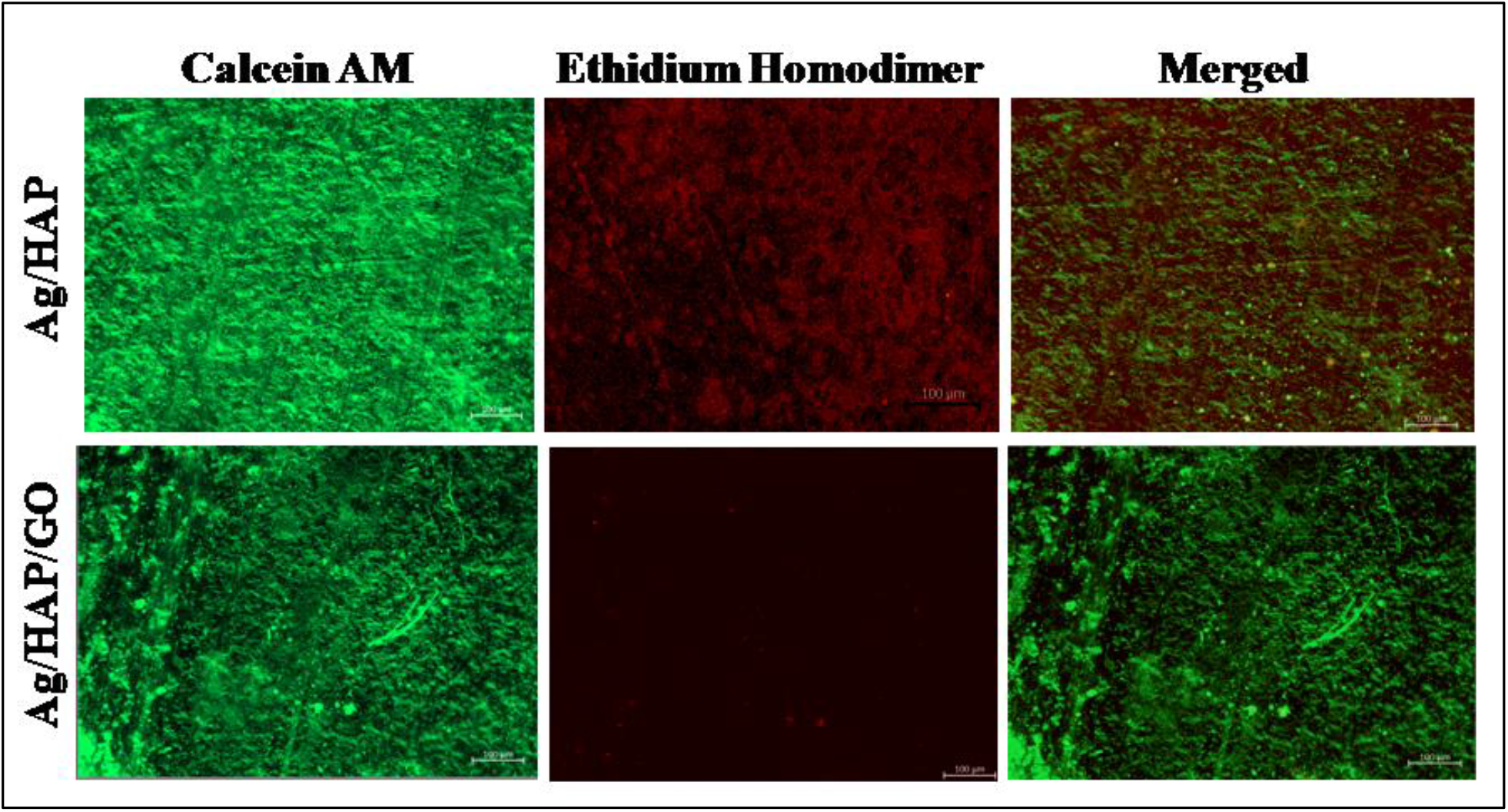
Live/dead imaging of hMSCs using double staining of Calcein AM and Ethidium Homodimer-1.

### 3.5. Alkaline phosphatase Activity

Alkaline phosphatase (ALP) is a common molecular marker for osteogenic differentiation, therefore ALP activity was a relevant test to assess the evolution of the induced osteogenesis in our experiment [17]. The cells seeded on agarose-hydroxyapatite-graphene oxide scaffolds for 7 days show significantly higher (p < 0.05) ALP activity than those seeded on agarose-hydroxyapatite scaffolds, respectively. These findings indicate that agarose-hydroxyapatite-graphene oxide scaffolds can stimulate the expression of early osteogenic proliferation markers without any external osteogenic agent. Earlier reports suggest that HAP and graphene oxide enhances the cellular ALP activity of osteoblastic cells [18]. The obtained results clearily suggests that HAP and graphene oxide work synergistically to enhance the differentiation of hMSCs.

### 3.6. Osteogenic markers expression

To examine the osteogenic differentiation of hMSCs on scaffolds, we analyzed the expression of early and late osteogenic markers by RT-PCR after 21 day of seeding. The expression of (ACAN), collagen-1A2 (COL1A2), Cartilage oligomeric matrix protein (COMP), collagen-1A1 (COL1A1), collagen-3A1 (COL3A1), bone morphogenetic protein-1 (BMP1), bone morphogenetic protein-6 (BMP6) and runt related transcription factor-2 (RUNX2) in hMSCs encapsulated scaffolds depicted an increasing trend (p < 0.05), as compared to control scaffolds. Therefore, the up-regulation of osteopontin and osteocalcin expression is correlated with RunX-2 expression in response to graphene oxide addition, which was a reflection of RUNX-2-mediated modulation of OPN and OCN (Figure 4, supplementary).

## 4. CONCLUSION

In summary, we have fabricated graphene oxide-reinforced agarose-hydroxyapatite 3D scaffolds by the additive methods, biocompatibility, and osteogenic potential were evaluated by in vitro studies. Comparatively, the graphene oxide reinforced scaffolds showed better biocompatibility and osteogenic potential than control scaffolds. Our results have suggested that graphene oxide reinforced scaffold could be a promising tool for the reconstruction of large bone defects, without using any exogenous cells or growth factors.

## ACKNOWLEDGMENTS

The authors acknowledge DST-New Delhi, India for financial support. The authors also acknowledge Centre for Nanosciences, IIT Kanpur, for providing Imaging and RT-PCR facility.

## Notes

### Competing Interest Statement

The authors have declared no competing interest.

